# Reusability Report: Few-shot learning creates predictive models of drug response that translate from high-throughput screens to individual patients

**DOI:** 10.1101/2023.07.06.547938

**Authors:** Emily So, Fengqing Yu, Bo Wang, Benjamin Haibe-Kains

## Abstract

Machine learning (ML) and artificial intelligence (AI) methods are increasingly used in personalized medicine, including precision oncology. Ma et al. (Nature Cancer 2021) developed a new method c alled “*Transfer of Cell Line Response Prediction*” (TCRP) to train predictors of drug response in cancer cell lines and optimize their performance in higher complex cancer model systems via few-shot learning. TCRP was presented as a successful modeling approach in multiple case studies. Given the importance of this approach to assist clinicians in their treatment decision process, we sought to reproduce independently the authors’ findings and improve the reusability of TCRP in new case studies, including validation in clinical trial datasets, a high bar for drug response prediction. Our results support the superiority of TCRP over established statistical and machine learning approaches in preclinical and clinical settings. We developed new resources to increase the reusability of the TCRP model for future improvements and validation studies.

## Introduction

With recent advances in molecular profiling and computational technologies, there has been an increasing interest in developing and using machine learning (ML) and artificial intelligence (AI) methods for personalized medicine and precision oncology. An active area of research focuses on computational models capable of predicting therapy response for cancer patients. Given the challenges present in obtaining a compendium of clinical genomic data that is sufficiently large for multivariable analysis, the majority of studies train drug response predictors from large-scale preclinical pharmacogenomic data and assess the performance of the resulting predictive models using limited patient data. Preclinical models, however, do not always faithfully recapitulate the therapy response observed in patients. Translating predictors trained on preclinical data to achieve good accuracy on clinical data remains an open challenge. If the translation is successful, these predictors hold great potential for improving the selection of anticancer therapies, a key challenge in precision oncology.

In a recent paper in Nature Cancer, Ma et. al introduced “*Transfer of Cell Line Response Prediction*” (TCRP)^1^, a new method for transfer drug response prediction based on few-shot learning. The authors showed that their approach enabled the development of computational models able to learn from immortalized cancer cell line data and predict response in more complex *in vitro* patient-derived cell cultures and *in vivo* patient-derived xenografts. Given the impressive nature of the original paper’s results, this Reusability Report aims to address two issues: (1) confirming the performance of the TCRP model in its published context; and (2) expanding its application on a larger compendium of preclinical pharmacogenomic and clinical trial data. Upon successful testing and deployment, modeling approaches such as TCRP, will help improve personalized medicine by facilitating the match of a patient molecular profile to optimal therapy.

## Reproducibility

We first aimed to fully reproduce the results outlined in the original publication: training drug response predictors using the GDSC1^2^ immortalized cancer cell line dataset and test its predictive value in patient-derived breast tumor cells (PDTC)^3^. This task was referred as ‘Challenge #2’ in the original publication, which is split into four stages: (1) extraction of gene expression and drug response features from the cell line dataset (GDSC1), (2) training the Model Agnostic Meta-Learning (MAML) algorithm to predict drug response in cell lines and hyperparameter tuning; (3) fine-tune these models using a small subset of the patient-derived tumor cells dataset (PDTC) using few-shot learning; and (4) validation on the remaining PDTC samples.

The authors shared the links to the GDSC1 and PDTC datasets, which are both publicly available. Their computer code was shared via two Github repositories, “[original codebase](https://github.com/idekerlab/TCRP)“ and “tcrp-reproduce“(both provided by the original authors). In the original codebase, a download link for the TCRP model is provided, with one line command to run the model on a single drug, Sorafenib. However, there is no instruction on model training or hyperparameter tuning on other drugs. In the “tcrp-reproduce” repository there are instructions for TCRP model training and baseline comparison to established predictive modeling approaches, however the instructions are incomplete. The software versions and dependencies are also listed.

In stage (1), we could only extract the input features (cell lines’ expression and mutational profiles, drug response values) for 32 out of 50 priority drugs investigated in the original publication due to missing reference files. Possible causes of this issue are discussed in Supplementary Information.

In stage (2), we were unable to reproduce the deep neural network (MAML) results exactly due to the missing drugs from stage (1) and the performance comparison was incomplete due to the lack of code for the baseline models’ implementation. Out of the four baseline models, only code for Random Forest, Linear Regression and K Nearest Neighbours was provided by the authors; the neural network was described as a “simplified version” of the TCRP model, and the architecture was not provided in the baseline code. Consequently, we had to reimplement the training and testing procedure of this important baseline model.

In stage (3), the hyperparameters for the fine-tuning process were not released with the original publication, forcing us to conduct a brute-force search to test all possible combinations of values listed by the authors in order to find the best-performing set of parameters for each drug-tissue combination. While the original authors provided methods for configuring the tuning for a subset of hyperparameters, we were still unable to get the exact objective values published which is most likely due to hyperparameter configurations we did not explore during cross-validation (**Figure 1**).

**Figure 1:**
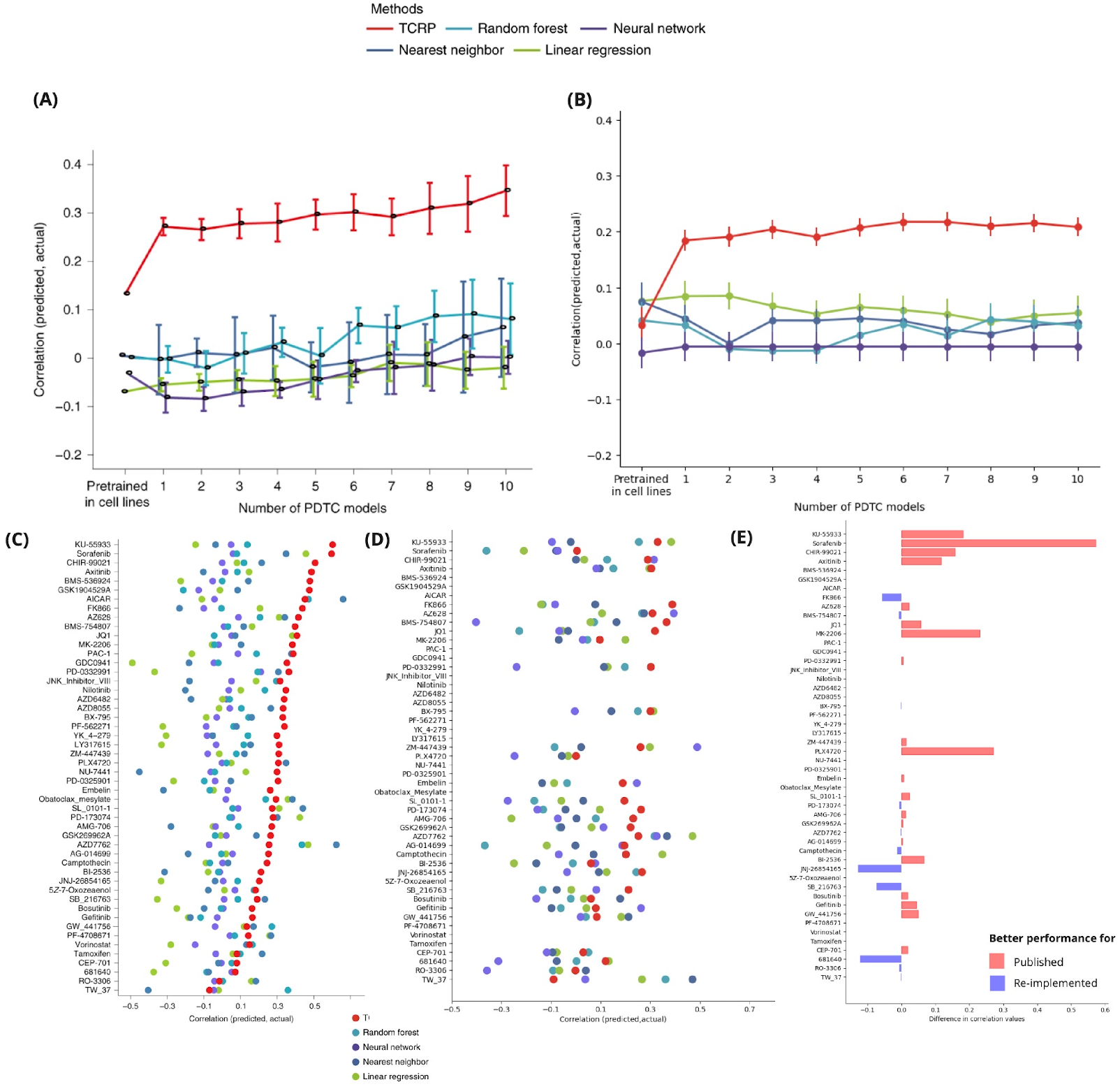
Reproducibility attempt of Challenge #2 from the original paper by Ma. et al 2021. (**A**) Original published results of Challenge 2. Results show that across 50 priority drugs, the TCRP model had higher Pearson’s correlation than baseline methods in predicting drug response in PDTCs when trained on GDSC1 cell lines (**B**) Our reproducibility attempt using resources and code provided in the original publication. The plot demonstrates that we were not able to reproduce the same objective value as published TCRP performance, but were able to prove it performs better than baselines **C)** Dot plot depicting original plot of published TCRP versus baseline performance, separated by drug. **D)** Dot plot depicting our reimplementation’s TCRP versus baseline performance, separated by drug. Once again there were differences in objective values of TCRP between our reimplementation and the original publication, but overall TCRP performance is better than baselines. **E)** Bidirectional plot showing the difference between the published objective value and the implementation objective value. Red bars (positive difference) signify the published model performed better for a drug’s prediction and blue bars (negative difference) signify the reimplemented model performed better for a drug’s prediction.

In stage (4), the authors showed that their TCRP model outperformed established statistical (Linear Regression) and machine learning models (Nearest Neighbor, Random Forest and Neural Network) on the set of 50 priority drugs on the PDTC dataset (**Figure 1A**). Using our reimplemented TCRP model, we were able to confirm the superiority of the TCRP model on average (**Figure 1B**). We did not achieve the same level of predictive value however, with Pearson correlation between predictions and measured drug response of over 0.3 in the original publication while the reimplemented model only reached a peak predictive value of 0.2 (**Figure 1B**). We also observed better predictive value for the simple linear regression model in our re-analysis, which reached positive Pearson correlation contrary to the published results (**Figure 1B**). At the drug level, there were consistent differences in our correlation values in comparison to the original publication (**Figure 1C**,**D)**. We observed the biggest differences for the drugs Sorafenib and MK-2206, which could be due to the lack of features for extraction (common genes between reference and transfer datasets), inconsistent names in drug targets or errors in the hyperparameter tuning (**Figure 1E)**.

Overall, we were able to reimplement the TCRP model and to confirm its superiority compared to established statistical and machine learning models using the same training (GDSC1) and validation (PDTC) datasets.

## Reusability

To assess whether the TCRP model can be applied to preclinical and clinical datasets that have not been explored in the original publication, we first investigated its performance when trained on different cell line datasets available on our ORCESTRA platform^4^, namely the CTRP^5^, gCSI^6^, GDSC2^7^ datasets. During the training process for each dataset, we again tuned the hyperparameter of the model for each drug. We observed that, for drugs shared among the different training sets, the hyperparameters used for optimal TCRP performance on one dataset can not seamlessly be used for another (**Supplementary Data 1**). Re-tuning hyperparameters for the TCRP model must occur anytime a new dataset (reference or validation) is introduced, or when different sets of drugs are used to extract features. The newly trained TCRP models still outperformed the baseline predictive models irrespective of the training set used (**Figure 2**), confirming the robustness of the approach with respect to cell line data. TCRP was the only model that was consistently the best across training datasets. Interestingly, for the only drug in common between all cell lines datasets – Paclitaxel – we observed that training TCRP on the CTRPv2 dataset achieved lower predictive value on PDTC than the reference datasets (Pearson of 0.48, 0.45 and 0.15 for the gCSI-, GDSC2-, CTRPv2-trained TCRP models, respectively), suggesting that the choice of training data may be important to develop the best drug response predictors.

**Figure 2:**
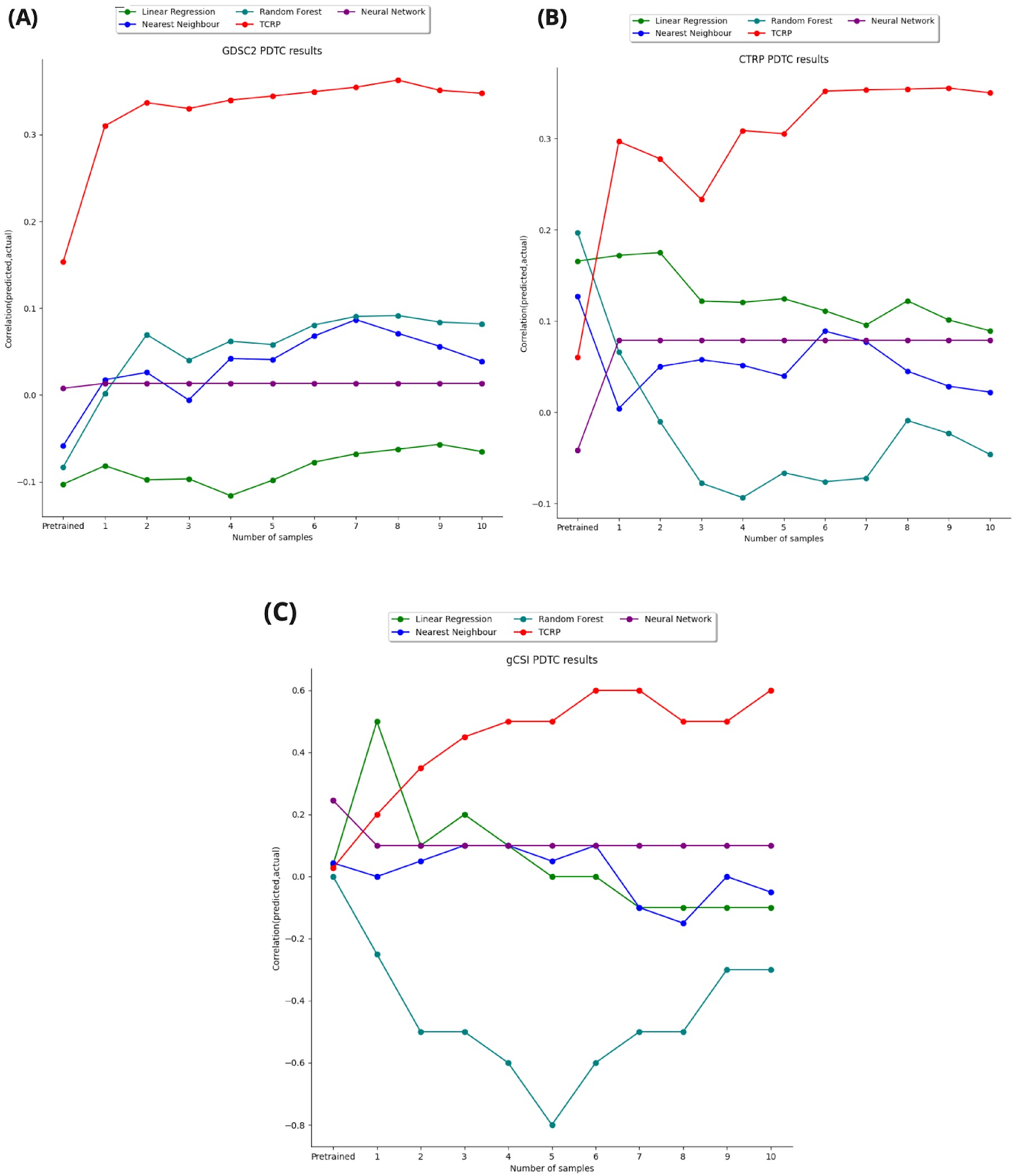
Performance of TCRP versus common baselines on new reference datasets. For each new reference dataset, instead of using the original 50 priority drugs, any drugs with sufficient number of cell line response values that were also present in the PDTC data were used. The drug sets are different across all 3 datasets. (**A**) performance of TCRP versus other baselines on predicting drug response in PDTCs, trained on gCSI (**B**) performance of TCRP versus other baselines on predicting drug response in PDTCs, trained on GDSC2 and (**C**)performance of TCRP versus other baselines on predicting drug response in PDTCs, trained on CTRPv2.

To further challenge the reusability of the TCRP approach, we assessed the predictive value of the TCRP model on datasets that are representative of diverse preclinical and clinical scenarios. Using GDSC1 as training set, we tested the transfer of predictors for response to paclitaxel in breast cancer (*i*) immortalized cell lines (UHNBreast^8,9^); (*ii*) patient-derived xenografts (Novartis PDX Encyclopedia^10^); and (*iii*) two clinical trials (GSE250066^11–13^ and GSE41998^14^). This is made challenging due to the fact that, in each setting, the drug response is assessed differently. For *in vitro* models, such as immortalized cancer cell lines or patient-derived tumor cell cultures, the drug response is measured via drug dose-response curves. For *in vivo* models, such as patient-derived xenografts, the drug response is measured via tumor-growth curves. In clinical trials, drug responses are evaluated using the Response Evaluation Criteria in Solid Tumors (RECIST)^15^, categorizing the tumor response as complete response (CR), partial response (PR), stable disease (SD), or progressive disease (PD). To achieve alignment between these different drug response readouts, it is possible to map all drug responses to categorical variables. For instance, Gao et al. introduce the modified RECIST (mRECIST) criterion to map tumor-growth curves to the RECIST categories used in clinical trials^10^. We^16^ and others^17^ have investigated ways to dichotomize drug response from dose-response curves in cell lines. To transfer drug response prediction across the *in vitro, in vivo* and clinical settings, we binarized the drug response readout as Responders (R) vs Non-Responders (NR). Specifically, we grouped CR and PR as R and SD and PD as NR. In cell-lines, we enforce a threshold cutoff on AAC (1-AUC), with AAC > 0.2 are classified as R, and AAC ≤ 0.2 as NR.

Training and validating TCRP in this new analysis framework, we observed that the TCRP model showed overall better performance than baseline models in most of the settings, including the two clinical trials (**Figure 3**). Notably, the TCRP model failed to outperform the nearest neighbor modeling approach in the UHNBreast cell line dataset (**Figure 3A**) while outperforming all other methods in the more complex patient-derived xenograft and clinical trial datasets (**Figure 3B-D**).

**Figure 3:**
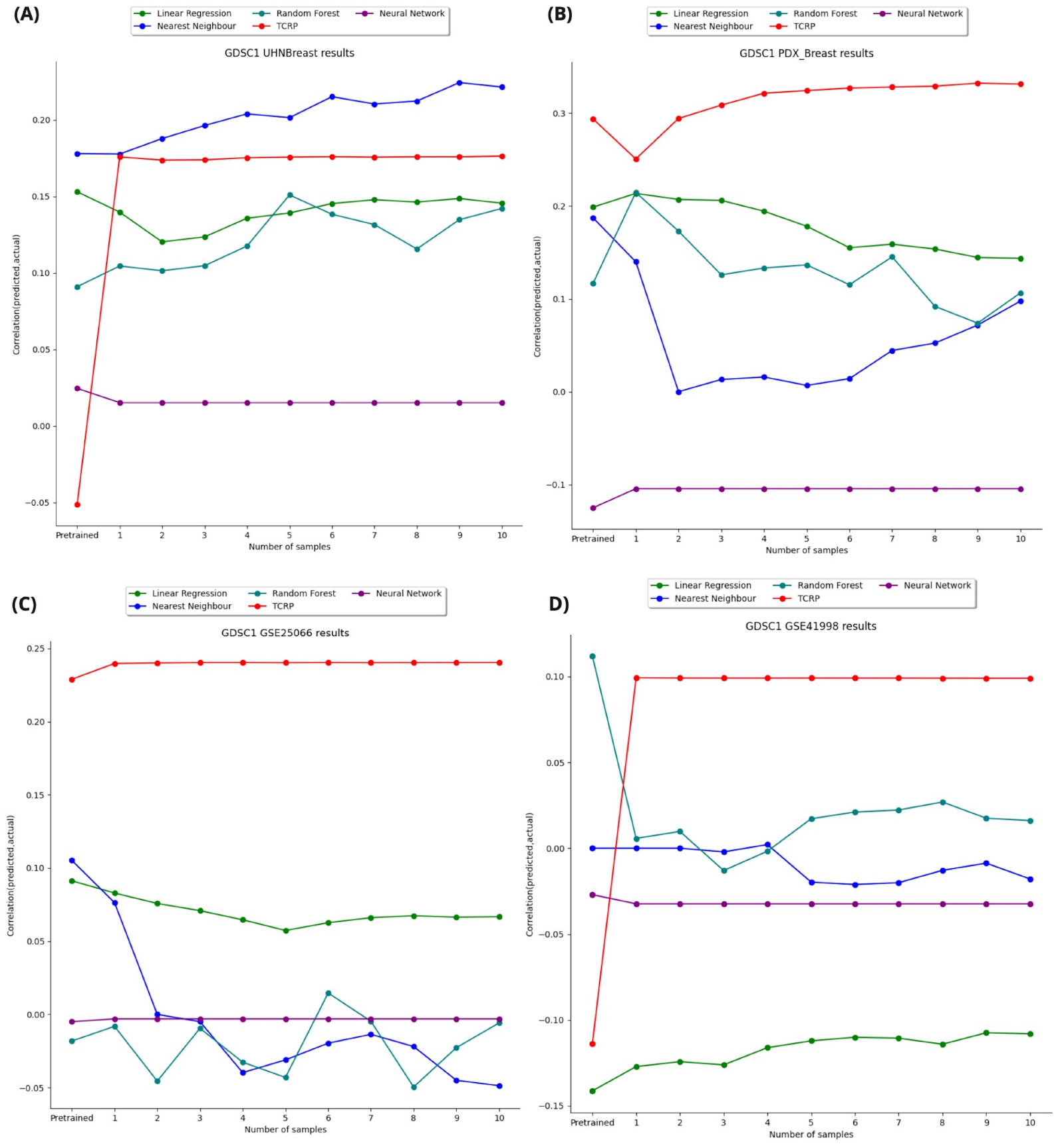
Performance of TCRP versus other baselines in three new validation contexts. Performance was evaluated using only drug response from Paclitaxel in all four datasets, and all four models were trained on GDSC1 (**A**) TCRP performance versus baselines on predicting drug response on new dataset of immortalized breast cancer cell lines (UHNBreast); (**B**) TCRP performance versus baselines on predicting drug response on breast cancer patient-derived xenografts models treated with Paclitaxel (Novartis PDX Encyclopedia) ; (**C**,**D**) TCRP performance versus baselines on predicting drug response on two clinical trials

Altogether our reusability study strongly supports the capacity of the TCRP modeling approach to transfer its predictive value by learning from few samples generated in a new context. Our results show that TCRP can learn the complex relationship between molecular features of cancer cells and their drug response using large cell lines’ pharmacogenomic datasets, and fine tune the resulting predictors to achieve higher performance in more complex preclinical models or in clinical trials.

## Discussion

In the course of reproducing and reusing the TCRP models, some persistent challenges were encountered. The primary issue is the limited number of drugs that we could recover from the reference datasets. Upon utilizing the same datasets as those employed in the original paper, only 32 out of 50 drugs were successfully recovered. Furthermore, upon re-applying the model to alternative reference datasets (i.e., baseline cell line drug screening studies), there was a substantial reduction in the number of drugs that the model was able to predict due to the smaller number of drugs commonly investigated.

One of the larger issues we faced in the reproducibility process was tuning the TCRP model for increased performance. The hyperparameters, including the number of few-shot samples and learning rate, must be uniquely adjusted for each specific drug and validation dataset combination. However, this process appeared to be worthwhile as it appeared that with tuning the TCRP model performed well in various contexts for reusability. There was consistent success in our experiments in predicting response in both cell line, patient-derived cancer models and clinical datasets.

Supporting the original publication, our results support the TCRP model as a successful approach to predicting clinical drug response. However, there are additional measures that can be taken to ensure the full reproducibility and reusability of the TCRP approach. Direct replication of a published model through code cloning and re-implementation can be a time-intensive and error-prone process, due to factors such as changes in path and directory and software environment requirements. Therefore, it is critical to ensure that the model is implemented in a manner that is easily shareable and executable^18^. To this end, all our experiments were conducted using a Code Ocean capsule, which offers an end-to-end workflow for the model and enables straightforward sharing and execution by other researchers^19^. Additionally, proper preprocessing of the data is crucial for the model’s application to other datasets. Our reference and testing datasets were sourced from ORCESTRA, which provides a curated dataset with detailed versioning and a unique digital object identifier (DOI), ensuring full transparency and reproducibility of dataset processing and subsequent analysis^4^. Through the integration and automation of the model construction process with data querying and processing, computational models such as TCRP can be readily evaluated and utilized by researchers, facilitating a greater impact of AI in medicine.

Our reproducibility and reusability study further cements the published few-shot learning model as a machine learning approach that could be applied to different contexts, including both cell line and clinical trial datasets. The success of TCRP in the clinical context suggests the great potential of this approach to improve precision oncology. In addition to introducing new contexts, there are improvements that can be made to the availability of this method for the scientific community. Methods such as the few-shot model can be presented in a containerized fashion, either in a python package or an interactive containerized repository on platforms such as Code Ocean.

## Data and Code Availability

The code to run and visualize the exact results of our reproducibility attempt as well as all our novel analyses for reusability can be found in this Code Ocean capsule (10.0260/CO.7288809.v2). Code used for analysis can also be found at github.com/bhklab/TCRP_Reusability_Report. All datasets used in our study are available on the data platform [ORCESTRA](https://www.orcestra.ca/).

## References

1. Ma, J. et al. Few-shot learning creates predictive models of drug response that translate from high-throughput screens to individual patients. Nat Cancer 2, 233–244 (2021).

2. Yang, W. et al. Genomics of Drug Sensitivity in Cancer (GDSC): a resource for therapeutic biomarker discovery in cancer cells. Nucleic Acids Res. 41, D955–61 (2013).

3. Bruna, A. et al. A Biobank of Breast Cancer Explants with Preserved Intra-tumor Heterogeneity to Screen Anticancer Compounds. Cell 167, 260–274.e22 (2016).

4. Mammoliti, A. et al. Orchestrating and sharing large multimodal data for transparent and reproducible research. Nat. Commun. 12, 5797 (2021).

5. Seashore-Ludlow, B. et al. Harnessing Connectivity in a Large-Scale Small-Molecule Sensitivity Dataset. Cancer Discovery vol. 5 1210–1223 Preprint at https://doi.org/10.1158/2159-8290.cd-15-0235 (2015).

6. Haverty, P. M. et al. Reproducible pharmacogenomic profiling of cancer cell line panels. Nature 533, 333–337 (2016).

7. Iorio, F. et al. A Landscape of Pharmacogenomic Interactions in Cancer. Cell 166, 740–754 (2016).

8. Safikhani, Z. et al. Gene isoforms as expression-based biomarkers predictive of drug response in vitro. Nat. Commun. 8, 1126 (2017).

9. Thu, K. L. et al. Disruption of the anaphase-promoting complex confers resistance to TTK inhibitors in triple-negative breast cancer. Proceedings of the National Academy of Sciences 115, E1570–E1577 (2018).

10. Gao, H. et al. High-throughput screening using patient-derived tumor xenografts to predict clinical trial drug response. Nat. Med. 21, 1318–1325 (2015).

11. Hatzis, C. et al. A genomic predictor of response and survival following taxane-anthracycline chemotherapy for invasive breast cancer. JAMA 305, 1873–1881 (2011).

12. Itoh, M. et al. Estrogen receptor (ER) mRNA expression and molecular subtype distribution in ER-negative/progesterone receptor-positive breast cancers. Breast Cancer Res. Treat. 143, 403–409 (2014).

13. Baldasici, O. et al. Circulating Small EVs miRNAs as Predictors of Pathological Response to Neo-Adjuvant Therapy in Breast Cancer Patients. Int. J. Mol. Sci. 23, (2022).

14. Horak, C. E. et al. Biomarker analysis of neoadjuvant doxorubicin/cyclophosphamide followed by ixabepilone or Paclitaxel in early-stage breast cancer. Clin. Cancer Res. 19, 1587–1595 (2013).

15. RECIST 1.1 – RECIST. https://recist.eortc.org/recist-1-1-2/.

16. Safikhani, Z., Smirnov, P., Freeman, M. & El-Hachem, N. Revisiting inconsistency in large pharmacogenomic studies. F1000Res 5, 2333. (2016).

17. Barretina, J. et al. The Cancer Cell Line Encyclopedia enables predictive modelling of anticancer drug sensitivity. Nature 483, 603–607 (2012).

18. Raff, E. Research Reproducibility as a Survival Analysis. AAAI 35, 469–478 (2021).

19. Clyburne-Sherin, A., Fei, X. & Green, S. A. Computational Reproducibility via Containers in Psychology. Mol. Pathol. 3, (2019).

