## Supplementary Information for "Reusability Report: Few-shot learning creates predictive models of drug response that translate from high-throughput screens to individual patients"

### Reproducibility of the original results

Stage (1): Extracting drug response data

The TCRP model needs first to be trained on large cell-line data, or the reference dataset, before transfer learning on PDTCs. In the data processing, we needed to find the common drug features between the 2 datasets. While we obtained the datasets using the provided link, after running the code for data processing, we only obtained the features for 32 out of 50 drugs in the paper. Further investigation shows that it is caused by the inconsistencies among the names of genes that the drugs are targeted. For example, “CDKs” were not matched with 'CDK9', 'CDK8', and thus the drug with mappable targets “CDKs” will be lost. The authors may reconcile the differences but with no clear indication on where and how they implement it. It would be helpful if there are reference files or instructions to convert between names and resolve the inconsistencies. Otherwise tracing each drug and gene names manually is required to recover more drugs.

When running the TCRP model on other datasets, the number of drugs that can be predicted was reduced. Upon further investigation, we discovered that the **assignment and distribution of training and testing samples was suboptimal, leading to instances where only a single testing sample was available. To ensure the accurate calculation of correlation coefficients, we manually removed these drugs with only one sample.**

Stage (2): Model Agnostic Meta-Learning

The implementation of the baseline model was absent from the pipeline. As the authors did not explicitly define the architecture for the CNN model, we had to implement it ourselves. We decided to follow the implementation of MAML model architecture, using one linear layer, ReLU activation layer, and batch normalization layer.

Stage (3): Fine-tuning via few-shot learning

The hyperparameters used for the fine-tuning process are as follows:
- Tissue number evolved in the inner update: (6, 12, 20)
- Meta learning rate and inner learning rate: (0.1, 0.01, 0.001)
- Number of hidden neurons: (5, 10, 15, 20)
- Number of layers: (1, 2)

For every drug, we trained the model using every possible combination of the hyperparameters with their values listed above. The hyperparameter set with the highest correlation is chosen for the specific drug model. It is computation-costly as the brute-force optimization method is done for each drug and each tissue, and for each time we trained on a new dataset.

Stage (4): Validation on independent data

With the selected hyperparameters in stage (3), we ran the drug-specific model and compared with the publication. We listed the drugs in the sequence of their predictive values as the original publication. The trend of the predictness of the drugs we identified mostly align with the original publication. Top predictive drugs include KU-55933, CHIR-99021. However, there are some drugs such as Sorafenib and MK-2206, which have surprising low correlation values in our re-implementation, despite being significantly higher values in the original publication.

### Supplementary Data: Hyperparameter values

The hyperparameter values used for prediction, along with all the objective values obtained for correlation, for predicted drugs can be found attached for all datasets. Different tables are labeled by reference and validation dataset.
